# The evolutionary potential of the influenza A virus hemagglutinin is highly constrained by intersegment epistasis

**DOI:** 10.1101/2022.05.19.492711

**Authors:** Tongyu Liu, Yiquan Wang, Timothy J C Tan, Nicholas C Wu, Christopher B Brooke

**Affiliations:** Department of Microbiology, University of Illinois at Urbana-Champaign, Urbana, Illinois, USA; Department of Biochemistry, University of Illinois at Urbana-Champaign, Urbana, Urbana, Illinois, USA; Center for Biophysics and Quantitative Biology, University of Illinois at Urbana-Champaign, Urbana, Illinois, USA; Carl R. Woese Institute for Genomic Biology, University of Illinois at Urbana-Champaign, Urbana, Illinois, USA; Carle Illinois College of Medicine, University of Illinois at Urbana-Champaign, Urbana, Illinois, USA

## Abstract

The ongoing antigenic evolution of the influenza A virus (IAV) hemagglutinin (HA) gene limits efforts to effectively control the spread of the virus in the human population through vaccination. The factors that influence and constrain the evolutionary potential of the HA gene remain poorly understood. Efforts to understand the mechanisms that govern HA antigenic evolution typically examine the HA gene in isolation and ignore the importance of balancing HA receptor-binding activities with the receptor-destroying activities of the viral neuraminidase (NA) for maintaining viral fitness. We hypothesized that the need to maintain functional balance with NA significantly constrains the evolutionary potential of the HA gene. We used deep mutational scanning to show that variation in NA activity significantly reshapes the HA fitness landscape by modulating the overall mutational robustness of the HA protein. Consistent with this, we observe that different NA backgrounds support the emergence of distinct repertoires of HA escape variants under neutralizing antibody pressure. Our results reveal a critical role for intersegment epistatic interactions in shaping the evolutionary potential of the HA gene.

## Introduction

Seasonal influenza A viruses (IAV) impose enormous public health (Paget et al., 2019) and economic burdens (Putri et al., 2018) across the globe on a yearly basis, despite the availability of licensed vaccines. The viral hemagglutinin (HA) glycoprotein mediates cell binding and entry and represents the primary target of protective neutralizing antibodies. The persistence of seasonal IAV lineages in the human population depends upon the continual accumulation of antigenically significant substitutions that facilitate evasion of humoral immunity elicited by prior infection or vaccination. This process, known colloquially as “antigenic drift”, necessitates yearly updating of seasonal influenza virus vaccines (Carrat and Flahault, 2007; Petrova and Russell, 2018; Yewdell, 2011).

The antigenic evolution of influenza viruses at the epidemiological scale appears highly constrained, as new antigenic variants typically only emerge every few years despite the high mutation rate of the virus (Brooke, 2017; Pauly et al., 2017; Petrova and Russell, 2018). Further, only a tiny fraction of potential escape variants generally emerge under immune selection, likely because most substitutions capable of reducing antibody binding avidity have deleterious pleotropic effects on HA function that offset their fitness benefits (Doud et al., 2017; Koel et al., 2013; Kosik et al., 2018). Defining the specific constraints that govern the emergence of antigenic variants is critical both for the design of next-generation vaccines that elicit more escape-resistant immune responses and for improving our ability to predict future evolutionary trends.

IAV encodes two primary surface glycoproteins: HA and neuraminidase (NA) that are irregularly distributed across the viral envelope (Harris et al., 2006; Michael Vahey and Fletcher Correspondence, 2019; Vahey and Fletcher, 2019; Wasilewski et al., 2012). HA and NA perform opposing functions during the viral life cycle: HA binds to sialic acid linkages to facilitate cell entry while NA typically cleaves sialic acid linkages to facilitate virion release (Kosik and Yewdell, 2019). Balancing these opposing functions is critical for maintaining viral fitness (Brooke et al., 2014; Gaymard et al., 2016; Kosik and Yewdell, 2019; Mitnaul et al., 2000; de Vries et al., 2020; Wagner et al., 2002; Xu et al., 2012a). Consistent with this, escape substitutions that emerge in HA are often associated with compensatory substitutions in NA that alter NA activity levels (Das et al., 2013; Hensley et al., 2011). While the importance of HA-NA balance is well-established, the implications of this critical functional relationship with NA for the evolutionary potential of HA have not been thoroughly examined. Here, we examine how phenotypic variation in NA can reshape the evolutionary landscape available to the HA gene, resulting in divergent pathways of antigenic escape.

## Results

### Changes in NA activity reshape the HA fitness landscape

Based on the importance of HA/NA functional balance for maintaining viral fitness, we hypothesized that phenotypic variation in NA would alter patterns of mutational tolerance of the HA protein. To test this, we generated two recombinant A/Puerto Rico/8/1934 (rPR8) viruses that were identical to wild type except for single amino acid substitutions in the NA protein that have been previously demonstrated to reduce NA activity. NA:K253R was first identified as a compensatory mutation that emerged following anti-HA antibody selection (Hensley et al., 2011), and NA:H274Y is a well-characterized NA inhibitor-resistance substitution (Bloom et al., 2010; Ives et al., 2002). As expected, NA:K253R reduced virion-associated NA activity ∼75% comparing with WT (**Fig 1A**). We further confirmed that NA:K253R reduced NA protein trafficking to the plasma membrane in infected Madin-Darby canine kidney (MDCK) cells (**Fig 1B**), and decreased the NA protein content of purified virions (**Fig 1C**). These data indicate that NA:K253R likely reduces virion-associated NA activity by reducing the amount of incorporated NA protein rather than by altering NA enzymatic activity. We failed to reproduce the previously reported adverse effects of NA:H274Y on NA activity (**Fig 1**). This inconsistency could be due to the different virus background used in this study. As our assays may fail to capture subtle but biologically significant effects of NA:H274Y on NA function in our system, we included it along with NA:K253R in subsequent experiments.

**Figure 1:**
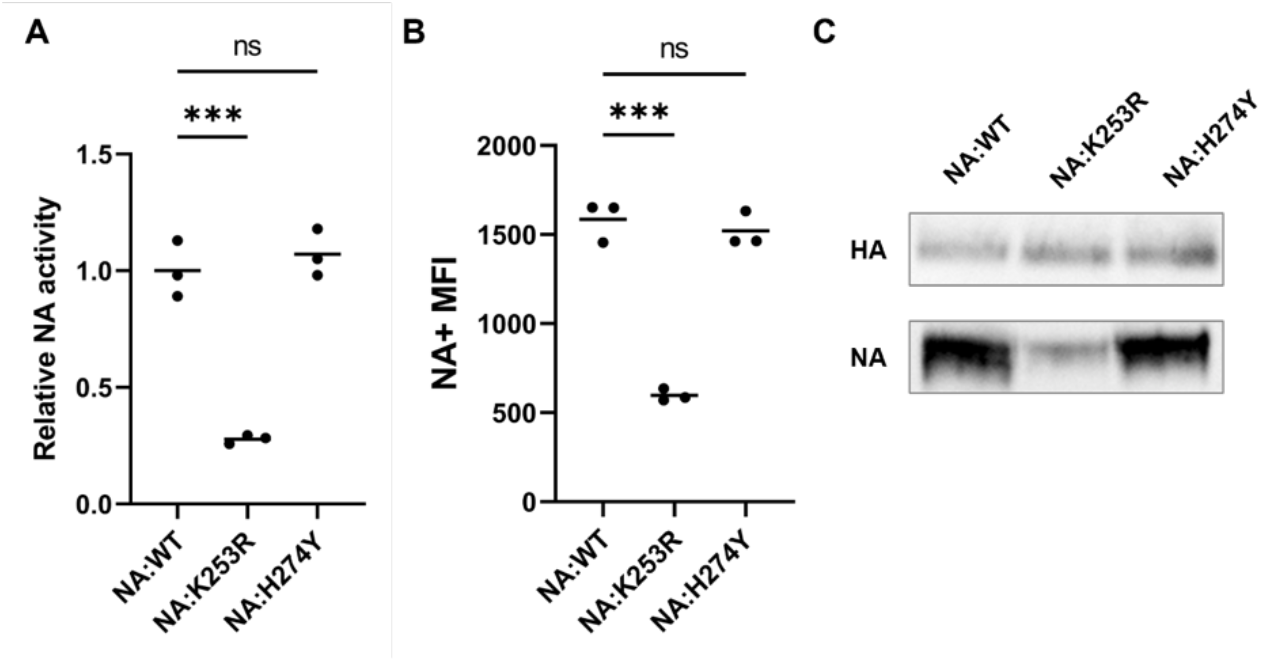
NA:K253R reduces NA surface expression and virion-associated NA activity. **(A)** The normalized V_max_ of NA activities from three independently generated virus stocks for each NA variant measured by MUNANA assay. The results were normalized by NP genome equivalents as determined by RT-qPCR. *** indicates p < 0.01 and ns indicates p > 0.05, based on t tests. **(B)** NA surface expression levels represented by mean fluorescence intensities (MFI) of NA positive cells at 16 hpi as measured by flow cytometry on MDCK cells infected at MOI = 0.05 TCID50/cell. *** indicates p < 0.01 and ns indicates p > 0.05, based on t tests. **(C)** Western blotting for HA and NA protein in purified virions. The input amounts of purified virions in the western blot were normalized based on the mean gray value of HA signal in a previous Western blot of the same samples.

We quantified the effects of NA:H274Y and NA:K253R on the mutational tolerance of the HA1 subunit of HA using deep mutational scanning (DMS) (Doud and Bloom, 2016; Fowler and Fields, 2014; Lee et al., 2018; Wu et al., 2014). We used a degenerate primer-based PCR approach to generate a reverse genetics plasmid library in which each codon in the HA1 domain was hyper-mutagenized to ensure sufficient representation of all possible amino acid substitutions as described previously (Doud and Bloom, 2016; Wang et al., 2021; Wu et al., 2014). We confirmed the presence of sufficient coding diversity within our plasmid library by deep sequencing **(Fig S1)**. For each NA genotype (WT, NA:H274Y, and NA:K253R), we rescued three independent recombinant virus populations encoding the mutagenized HA1 domain (HA1dms) and a WT PR8 backbone using the established IAV reverse genetics transfection system. We passaged each population once in MDCK cells for 24 hours at a starting multiplicity of infection (MOI) of 0.05 TCID50/cell to minimize cellular co-infection and thus maintain genotype-phenotype linkages. We then performed barcoded sub-amplicon deep sequencing on each post-passage virus population, along with the mutagenized HA plasmid library used to generate the viruses.

For every possible amino acid substitution in HA1, we calculated an enrichment factor by dividing its post-passage frequency by its frequency in the input plasmid library. We then calculated normalized relative fitness scores for each missense substitution in HA1 by normalizing based on the enrichment factor distributions of nonsense and synonymous substitutions. In brief, we assumed all nonsense substitutions would be lethal, regardless of genetic background and set their mean relative fitness values at 0 for each experimental replicate. Similarly, we assumed all synonymous substitutions would be neutral and set their mean relative fitness values at 1 for each experimental replicate. The pairwise correlation coefficients of fitness scores for specific substitutions between experimental replicates ranged from 0.665 to 0.812, indicating that our fitness effect measurements were highly reproducible (**Fig 2A, S2**).

**Figure 2:**
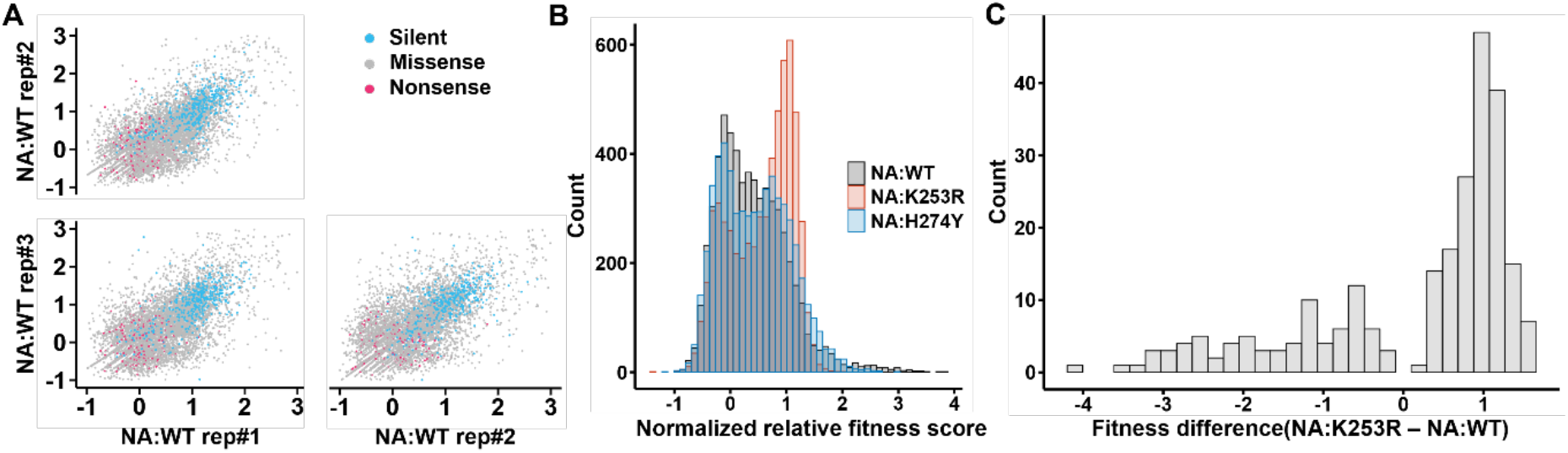
Variation in NA backgrounds affects the global fitness landscape of HA. **(A)** Correlations of normalized relative fitness scores across NA:WT replicate populations. Each dot represents the normalized relative fitness score of a specific substitution in the two indicated samples. Silent, missense, and nonsense substitutions are colored as indicated in the legend. **(B)** The distributions of normalized relative fitness scores of all missense substitutions in the indicated genetic backgrounds. Values are averaged across three replicates for each genotype. **(C)** The distribution of differences in missense substitution fitness scores between NA:K253R and NA:WT. Only shows substitutions for which t test on differences between genotypes yielded p < 0.01.

A large majority of missense substitutions in the rPR8-WT background had normalized fitness scores of <1, with the overall peak of fitness effects near 0, while only a tiny minority had fitness scores >1, as expected given that this virus is fairly well adapted to this host system (**Fig 2B**). This distribution is also consistent with previous studies across multiple virus families (including IAV) that indicate the vast majority of mutations have deleterious effects on relative fitness (Sanjuan, 2010; Sanjuan et al., 2004; Visher et al., 2016; Wu et al., 2014). Surprisingly, the normalized distribution of fitness effects (DFE) for rPR8-NA:K253R was shifted significantly compared with WT (**Fig 2B,C**), with a smaller peak of lethal or near-lethal substitutions with fitness scores of ∼0 and a large increase in substitutions with fitness scores of ∼1, indicating neutral or nearly-neutral effects on relative fitness. Substitutions at 150 out of 325 HA1 residues in rPR8-NA:K253R exhibited significant shifts (p<0.01, t test) in fitness scores compared with rPR8-WT. Finally, the DFE for rPR8-NA:H274Y was also shifted but to a lesser extent, consistent with the minimal effect of this substitution on NA function (**Fig S3**). Altogether, these data suggest that phenotypic variation in NA can have widely distributed effects on the mutational tolerance of the HA gene.

### Epistatic effects of NA on HA mutational fitness effects are enriched near the HA receptor binding site

To better understand how phenotypic variation in NA can affect mutational tolerance at specific residues, we calculated the differences in normalized fitness scores of individual HA1 substitutions between rPR8-WT and the two NA variants (**Fig 3A, S4**). In this analysis, positive difference values for individual substitutions indicate higher fitness in the NA variant background relative to WT while negative values indicate a higher fitness in the rPR8-WT background. Negative fitness score differentials were clearly enriched in a subset of residues, many of which are located near the receptor binding site (RBS).

**Figure 3:**
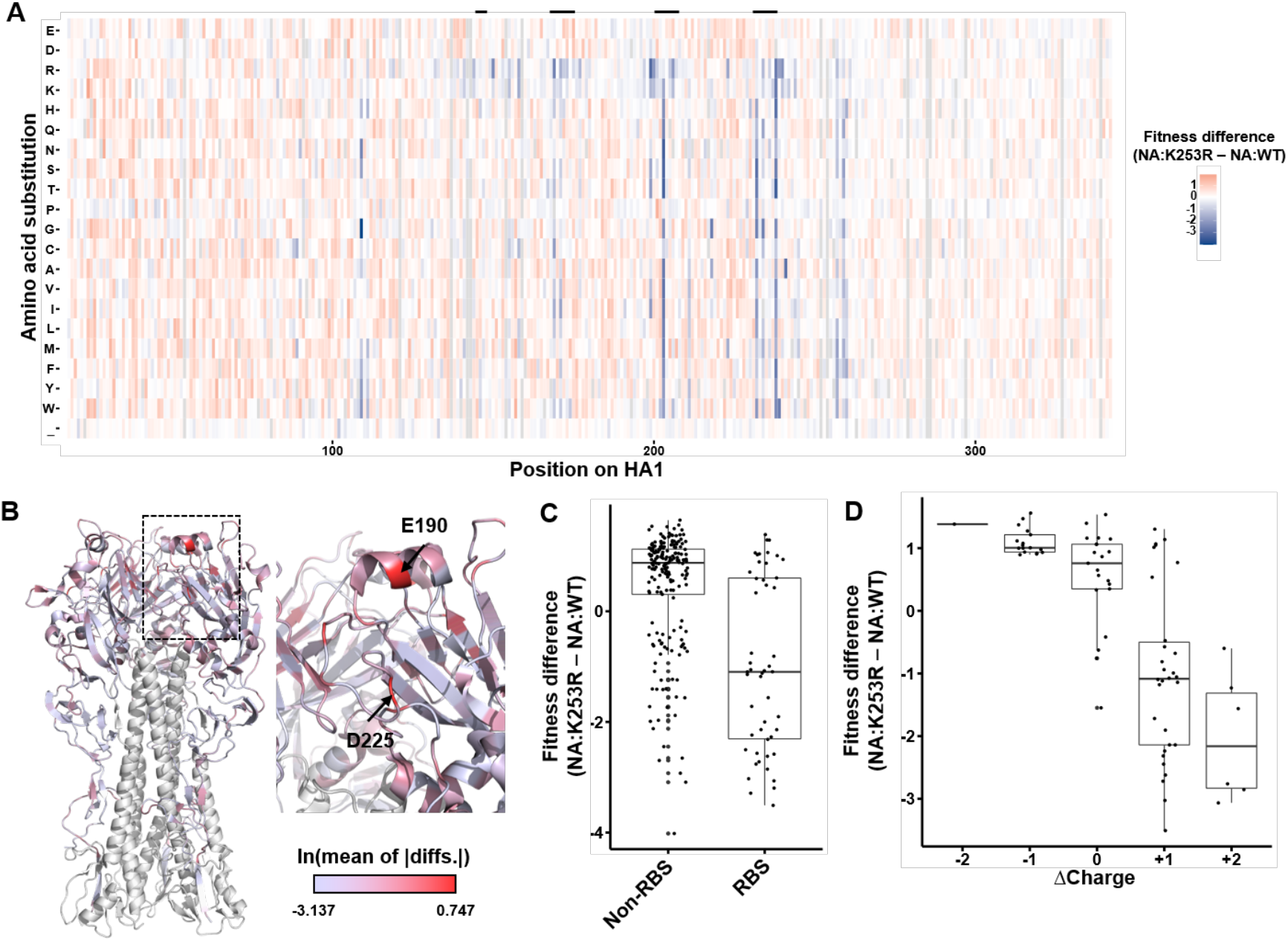
Epistatic effects of NA on HA mutational tolerance are concentrated in residues associated with receptor binding. **(A)** Normalized relative fitness score differences between NA:WT and NA:K253R for each substitution at each residue in HA1 as measured through DMS. Each value was generated by subtracting the mean fitness score of 3 replicates for each genotype. Gray indicates that the mutation had insufficient coverage in the plasmid library. Residue numbering based on the initiating methionine. Secondary structures forming the receptor binding site (130 loop, 150 loop, 190 helix, 220 loop) are indicated by the black bars above. **(B)** HA structure (PDB:1RU7) showing all HA1 residues colored by the natural log values of per-residue mean of absolute differences (MAD) of all substitutions between NA:WT and NA:K253R. HA2 domain colored in white. **(C)** Normalized relative fitness score differences between residues in HA1 associated with the receptor binding site versus those that are not (only showing substitutions with p < 0.01 by t test for comparison between NA:WT and NA:K253R). **(D)** Correlation between normalized relative fitness score differences and charge changes on surface (only showing substitutions with p < 0.01 by t test for comparison between NA:WT and NA:K253R).

Since rPR8-NA:K253R and rPR8-WT exhibited the most profound difference in NA activity, we focused on them for the following analyses. To define how NA influences mutational tolerance across the HA structure, we calculated per-residue mean of absolute difference (MAD) values (quantifies effects on overall mutational tolerance) of every residue in HA1 and plotted these values on the HA structure (**Fig 3B**). Key residues involved in receptor specificity, including E190 and D225, showed dramatically higher MAD values, indicating that epistatic interactions with NA are enriched in residues involved in receptor binding. Overall, NA-dependent fitness differences were significantly lower for residues associated with the RBS compared with those elsewhere in HA1 (p = 0.003739, unpaired two-samples Wilcoxon test) (**Fig 3C**). In other words, substitutions in the RBS tended to have a higher fitness in rPR8-WT than rPR8-NA:K253R, whereas the substitutions in HA1 non-RBS regions showed the reverse.

From the heatmap in **Fig 3A**, we observed that the positively charged amino acids arginine and lysine exhibited distinct mutational fitness effect patterns from other amino acids. We hypothesized that substitutions that result in net positive charge changes may be more tolerated within the WT NA background as these changes would likely enhance binding to the negatively charged cell surface via electrostatic intercations. As expected, negative charge changes on HA surface determined by GetArea (Fraczkiewicz and Braun) were more tolerated in rPR8-NA:K253R while positive charge changes had higher fitness in rPR8-WT (**Fig 3D**). Altogether, these observations indicate that the epistatic effects of NA phenotype on HA mutational tolerance are most pronounced for residues involved in receptor binding and cell adhesion.

### HA takes distinct mutational pathways to escape neutralizing antibody pressure depending on NA genotype

Our DMS data clearly demonstrated that variation in NA activity can significantly alter the mutational tolerance of the HA1 domain, particularly at residues surrounding the RBS that are known to be highly antigenically significant (Caton et al., 1982; Koel et al., 2013). Based on this, we hypothesized that different NA backgrounds would support the emergence of distinct repertoires of escape variants under neutralizing antibody selection. To test this, we performed *in vitro* selection experiments using the Sb epitope-specific neutralizing monoclonal antibody (mAb) H36-26 and the three recombinant viruses detailed above (rPR8-WT, rPR8-NA:K253R, and rPR8-NA:H274Y). Importantly, all three viruses have identical HA sequences. To minimize variation introduced by bottlenecking during virus rescue, we pooled three independent rescues of each virus to generate the parental virus populations for selection experiments. We infected MDCK cells with 10^7^ TCID50 of each virus in sextuplicate in the presence of H36-26 at a concentration where neutralization is saturated under these conditions (**Fig S4**). We passaged viral populations twice (16 hours each passage) in the presence of H36-26 to reach sufficient titers for sequencing (>10^4^ TCID50/mL). We then deep sequenced post-selection viral populations and identified single nucleotide variants (SNVs) that emerged above background using DeepSNV (Gerstung et al., 2012, 2014).

Distinct repertoires of escape substitutions emerged in mutant and WT NA backgrounds. In the WT background, HA:E156K emerged to high frequency (>60%) in 6/6 replicate populations, while K189Q was also observed at frequencies between 2% and 20% in 5/6 populations (**Fig 4A**). In contrast, neither E156K nor K189Q were observed above background in either rPR8-NA:K253R or rPR8-NA:H274Y. Instead, Q196K/R substitutions emerged to high frequency in 5/5 (one of the replicates in rPR8-NA:K253R failed to grow) or 6/6 replicates for rPR8-NA:K253R and rPR8-NA:H274Y, respectively. Q196R was present above background in 2/6 WT populations while Q196K was not observed. Both E156 and Q189 are located within the canonical Sb epitope (**Fig 4B**). These results are consistent with the DMS data, where HA:E156K showed higher relative fitness than HA:Q196K in the rPR8-WT background while HA:Q196R showed higher relative fitness than HA:E156K in the rPR8-NA:K253R background (**Fig S6**).

**Figure 4:**
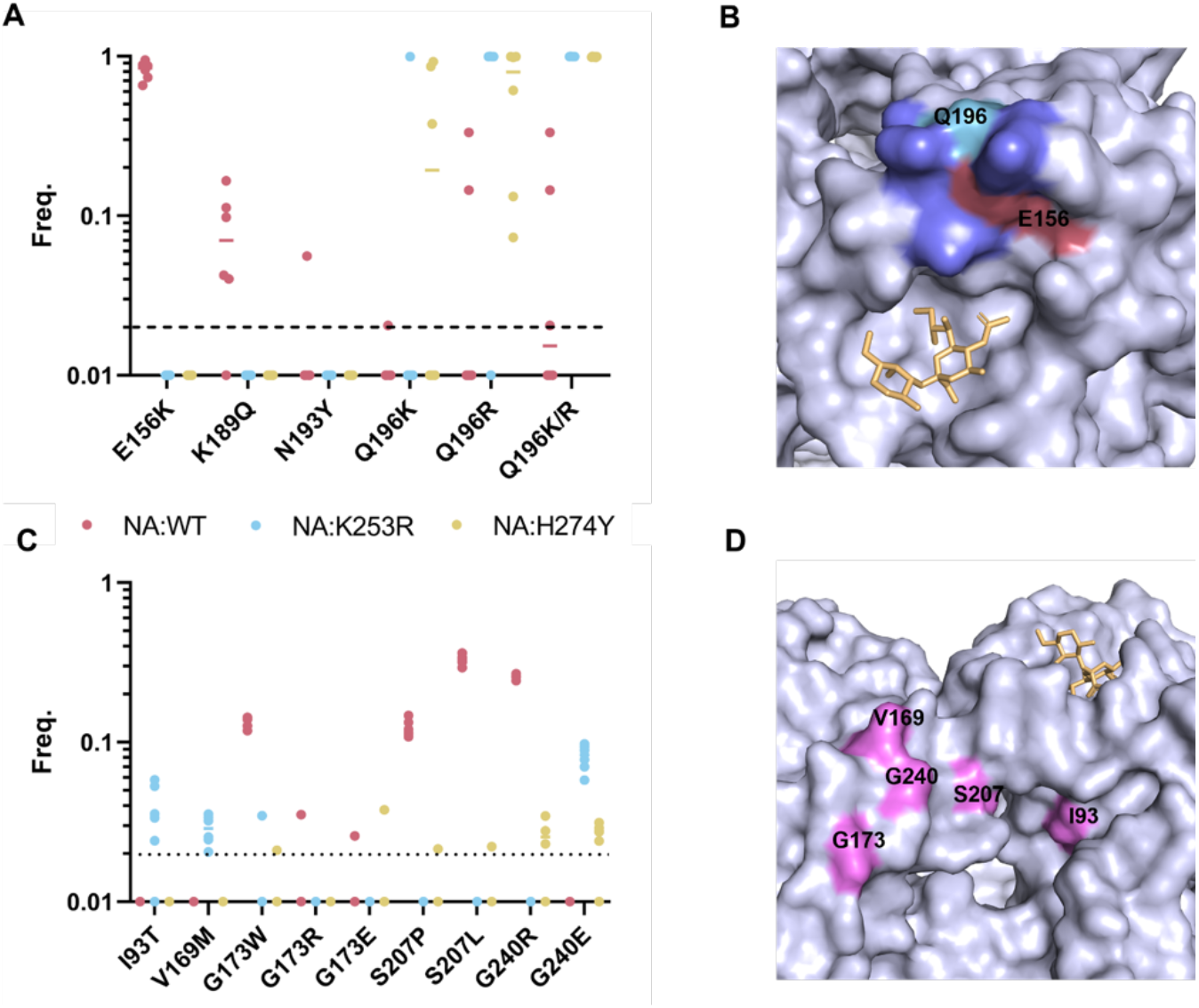
Different NA backgrounds support distinct mutational pathways to escape from neutralizing anti-HA antibodies. **(A)** Six independent populations of PR8-WT, PR8-NA:K239R, and PR8-NA:H274Y were passaged twice in MDCK cells in the presence of a neutralizing concentration of the anti-HA mAb H36-26, deep sequenced and analyzed via deepSNV(Gerstung et al., 2012). Frequencies of all HA amino acid substitutions detected at frequencies above the 2% frequency threshold (dashed line) across all replicates. Data from no mAb controls not shown but had no variants above 2%. Each dot represents the frequency in a single replicate. **(B)** The position of sialic acid receptor (yellow), E156 (rose), Q196 (cyan), and other Sb epitope residues (Caton et al., 1982) (purple) on the HA structure (PDB: 3UBQ (Xu et al., 2012b)). **(C)** The frequency of all HA amino acid substitutions detected under H17-L2 selection. Same experimental design as in (A) but using the Ca1-specific mAb H17L2. No mAb controls not shown but had no variants above 2% frequency (indicated by the dashed line). **(D)** The position of H17-L2 escape substitutions and the canonical Ca1 epitope in pink and sialic acid receptor (yellow) (Caton et al., 1982) on HA structure (PDB:3UBQ (Xu et al., 2012b)).

We hypothesized that viruses with lower relative NA activity would support the emergence of HA escape variants with lower receptor binding avidity compared with viruses with higher NA activity. We compared the receptor binding avidities of HA:E156K (predominant variant that emerged exclusively in the rPR8-WT background) and HA:Q196K (predominant variant that emerged exclusively in the NA mutant backgrounds) using bio-layer interferometry (BLI). As expected (Hensley et al., 2009), HA:E156K exhibited a higher receptor binding on-rate compared with HA:Q169K (**Fig S7**).

To test the generality of this observation, we performed a similar selection experiment using the Ca1-specific mAb H17-L2. Again, we observed distinct repertoires of escape variants in WT versus NA mutant backgrounds (**Fig 4C**). G173W, S207P/L and G240R were the dominant substitutions that emerged above background for rPR8-NA:WT viruses but were only sporadically observed above background in the rPR8-NA:K253R and rPR8-NA:H274Y backgrounds. Instead, I93T, V169M, and G240E were exclusively identified within the rPR8-NA:K253R and/or rPR8-NA:H274Y backgrounds. All these residues are located within or adjacent to the canonical Ca1 epitope (**Fig 4D**). Our results clearly demonstrate that different NA genotypes can reproducibly foster the emergence of distinct repertoires of HA escape variants under neutralizing antibody selection.

## Discussion

The specific factors that govern the evolutionary potential of the IAV HA gene remain poorly understood. Our results reveal how phenotypic variation in NA can profoundly reshape the fitness landscape available to the HA gene, thus determining its potential for future adaptation. The importance of balancing the opposing activities of the HA and NA glycoproteins for maximizing viral fitness is well established (Kosik and Yewdell, 2019). The importance of epistatic networks within HA has also been demonstrated to influence antigenic evolution (Kryazhimskiy et al., 2011; Wu et al., 2018). Here, we extend these concepts by demonstrating how intersegment epistasis arising from the intimate functional relationship between HA and NA significantly constrains the evolutionary potential of HA.

Given the typical number of contact residues involved in neutralizing antibody binding, numerous substitutions across multiple residues could potentially mediate escape. Most of these potential escape substitutions have deleterious pleiotropic effects on HA function that severely limit their overall viability and emergence potential (Doud et al., 2017; Kosik et al., 2018; Wu and Wilson, 2017). Our data indicate that as the HA genes of seasonal influenza viruses evolve to escape humoral immune pressure, the specific mutational pathways taken will be highly contingent upon the associated NA gene. Given that multiple N1 and N2 clades are often co-circulating, our results strongly suggest that the evolution of the HA gene cannot be viewed in isolation and that efforts to predict the evolutionary trajectories of seasonal IAVs must account for the influence of the associated NA segment.

Based on our overall hypothesis, we expected to observe a strong epistatic relationship between NA and the HA residues involved in receptor binding. Specifically, we expected substitutions that increase HA receptor binding avidity to have higher relative fitness in the context of NA:WT compared with NA:K253R, while substitutions that reduce receptor binding avidity to have higher relative fitness in the context of NA:K253R. Consistent with this, we observed that the fitness effects of substitutions associated with the RBS were highly sensitive to the NA background. This pattern was also observed in the context of neutralizing mAb escape, where HA:E156K, an escape substitution that significantly increases receptor binding avidity, dominated in the context of NA:WT but never emerged above background in NA:K253R. Instead, NA:K253R supported the emergence of an alternative escape substitution, HA:Q196K, that was associated with a less pronounced increase in receptor avidity.

Surprisingly, we discovered that the mutational tolerance profiles of numerous residues distal from the RBS were also significantly affected by the NA background. Epistatic effects of NA on these residues may still be largely driven by HA/NA balance issues, however, as numerous mechanisms can influence the ability of HA to facilitate receptor binding. These effects could involve surface charge changes that modulate electrostatic interactions between virions and the negatively charged cell surface. Another possibility is that destabilizing substitutions in HA (that potentially decrease proper folding, trafficking, virion incorporation, and/or receptor binding) may be better tolerated in genetic backgrounds with lower relative NA activity.

Our data demonstrate how variation in NA activity can modulate the overall mutational robustness of HA beyond the cluster of residues involved in receptor binding. We found that NA:K253R was associated with a positive overall shift in the mean fitness effects distribution for HA1, indicating that the relative fitness costs of a large number of substitutions were decreased in the context of reduced NA activity. We hypothesize that this effect is due to the functionally opposed primary activities of HA and NA. Viruses with lower relative levels of NA activity will have weaker functional constraints on HA, as they will better tolerate decreases in HA receptor binding avidity associated with amino acid substitutions. In this way, the functional balance of HA and NA may provide a simple system for studying how intergenic epistatic interactions can influence the mutational robustness of a viral protein.

The variability in the overall mutational robustness of the HA gene as a function of NA phenotype has substantial broader implications for IAV evolution. Mutational robustness has been hypothesized to facilitate the adaptive potential of proteins (and by extension, viral populations) under some conditions (Bloom et al., 2006; Draghi et al., 2010; Elena, 2012; Lauring et al., 2013; McBride et al., 2008; de Visser et al., 2003). By buffering mutational fitness effects, increases in robustness can promote the accumulation of genetic variants that may confer enhanced fitness or rescue in changing environmental conditions. In an extreme example, a zoonotic IAV population encoding a more mutationally robust HA gene would be more likely to accumulate substitutions that could facilitate successful cross-species transmission. Alternatively, a recent study demonstrated that increasing the robustness of a bacteriophage protein by increasing its thermostability actually decreased the potential of the virus to evolve to expand its host range (Strobel et al., 2022). In the context of IAV antigenic evolution, deleterious mutation load has been hypothesized to govern the potential for antigenic escape variants to emerge at the host population level, suggesting another mechanism by which variation in HA mutational robustness could influence antigenic drift (Koelle and Rasmussen, 2015). Altogether, our results suggest that variation in NA may have profound, if difficult to predict, effects on the evolvability of the HA gene.

In conclusion, our results demonstrate that epistatic interactions between HA and NA play a major role in shaping the fitness landscape of the HA gene (and almost certainly the NA gene as well) and in determining the most likely genetic pathways of antigenic evolution. Thus, the need to maintain functional balance between HA and NA activities imposes a significant constraint on the evolutionary potential of influenza viruses and should be considered in efforts to predict future evolutionary trjectories.

## Methods

### Cells and viruses

Madin-Darby canine kidney (MDCK) and human embryonic kidney HEK293T (293T) cells were passaged and maintained in Minimum Essential Medium (MEM + GlutaMAX, ThermoFisher Scientific) with 8.3% fetal bovine serum (FBS, Avantor Seradigm Premium Grade Fetal Bovine Serum) in 37°C and 5% CO2. MDCK cells and 293T cells were gifts of Dr. Jonathan Yewdell and Dr. Joanna Shisler, respectively.

A/Puerto Rico/8/1934 (PR8) and the specific PR8 mutants (generated by PCR mutagenesis) used in the study were generated via standard reverse genetics. In brief, ∼60% confluent 293T cells in 6-well plate were transfected with 500 ng of each segment cloned in the pDZ reverse genetics vector (jetPRIME, Polyplus Transfection). After 24 hours, the medium was replaced by the infection medium (MEM + 1 μg/mL TPCK-treated trypsin + 1 mM HEPES and 50 μg/mL gentamicin). Supernatants from transfected cells were collected at 48 hours post transfection and used to infect MDCK cells in 6-well plate to generate the seed stock. Seed stocks were collected at 48 hours post infection (hpi) or upon development of cytopathic effect (CPE), which ever came first. Working stocks were generated by infecting MDCK cells in T75 or T175 flask with seed stock at an MOI of 0.0001 TCID50/cell and collecting the supernatant at 48 hpi or when early signs of CPE were observed, which ever came first. Virus stocks were tittered by standard TCID50 assay. The plasmids were generously provided by Dr. Jonathan Yewdell.

The secondary structures forming the receptor binding site are defined as: T132-V135 (130 loop), T155-P162 (150 loop), N187-Y195 (190 helix), A218-D225 (220 loop) (Tzarum et al., 2017; Wu et al., 2013). The residue numbering of HA and NA is based on alignment to structural numbering of H3N2 (strain A/Hong Kong/1/1968 H3N2 (Brown et al., 2001), UniProt: Q91MA7, Q91MA2 for HA and NA) unless specified otherwise.

### MUNANA assay

The substrate 2′-(4-Methylumbelliferyl)-α-D-N-acetylneuraminic acid sodium salt hydrate (MUNANA, Sigma-Aldrich) was dissolve in NA buffer (33 mM MES, 4 mM CaCl2 within 1X PBS, pH = 6.5) and aliquoted. A black 96-well half-well flat bottom plate, the plate reader, NA buffer and the substrate was preheated to 37°C. 25 μL of virus sample (diluted in NA buffer) was mixed with 20 μL of the substrate (200 μM) and taken to the plate reader to measure the fluorescent kinetic (excitation wavelength = 365nm, emission wavelength = 450 nm) for 45 min. V_max_ values were estimated based on data collected after the first 10 min of the assay. The results were normalized based on the genome equivalent of NP segment determined by RT-qPCR.

### Cellular surface staining of NA

MDCK cells were infected with MOI = 0.05 based on the TCID50 titer of the viruses. MEM + 8.3%FBS was added on the cells after infection for 1 hour and replaced by NH_4_Cl medium (MEM, 50 mM HEPES, 20 mM NH_4_Cl, pH = 7.2) to block secondary infection. Cells were collected 16 hpi and stained with NA antibody (NA2-1C1-AF488, 1:1600) without permeabilization and run on a BD FACSAria Flow Cytometer. The NA positive cells were gated based on the no infection controls and the expression level were measured by the mean fluorescence intensity.

### Western blot of HA and NA

The virus stocks used for western blot were purified by ultra-centrifugation. Briefly, 10 mL of 20% sucrose with 30 mL of the virus stock was centrifuged at 27000 rpm, 4°C for 2 hours. The pellet was dissolved in PBS overnight. The product was then added on the top of a cushion of 15% sucrose and 60% sucrose and centrifuged. The virus band was collected after and washed by PBS. The purified viruses were heated at 98°C for 2 min, loaded on Bis-Tris protein gel (Bolt 4-12% Bis-Tris Plus, invitrogen), run on 150 V for 45 min and then transferred to PVDF membrane (iBlot2 PVDF Mini Stacks, invitrogen). The membrane was then stained with HA antibody (RA5-22, obtained through BEI Resources) and rabbit anti-NA polyclonal antibody (gift of Dr. Jonathan Yewdell).

### Deep mutation scanning of HA1

To generate all possible amino acid substitutions within HA1, NNK was introduced into each codon of the interest (D18-S342, H1 numbering from the first amino acid residue) by overlapping PCR (Phusion High-Fidelity DNA Polymerase, ThermoFisher Scientific) to generate full-length HA amplicons. NNK-mutagenized HA amplicons were then cloned into pDZ vector by T4 ligation (T4 DNA Ligase, New England BioLabs Inc.). The ligation product was then used to transform DH10B competent cells (MegaX DH10B T1, Invitrogen) and yield 9.8×10^5^ colonies (150X coverage of all substitutions) in total from two transformation reaction. The colonies were harvested and the plasmids were extracted by Midi prep (HiSpeed Plasmid Midi Kit, QIAGEN).

To rescue the viruses, 1.25×10^7^ 293T cells and 6×10^6^ MDCK cells were mixed and seeded per T175 flask in 25 mL of cell growth medium (MEM + 8.3% FBS). The next day, 7 μg each of the 8 reverse genetics plasmids was mixed with 112 μL of jetPRIME reagent (Polyplus Transfection) and 1.2 mL jetPRIME buffer (Polyplus Transfection) and this mixture was added to the medium. After 24 hours of transfection, the medium was removed, cells were washed with PBS, and 20 mL of infection medium was added. Supernatants were collected 72 hours post transfection. Each virus was passaged for 16 hours in present of same concentration of the antibody once in a T175 flask of confluent MDCK cells at a starting MOI of 0.05 TCID50/cell.

Viral RNA was extracted (QIAamp Viral RNA Kits, QIAGEN) and used to generate the cDNA (SuperScript III, ThermoFisher Scientific). For RT reaction, 11.8 μL of viral RNA was mixed with 1 μL random hexamer (125 ng/μL) and incubated at 65°C for 5 min for primer binding. The mixture was then added to be a 20 μL reaction and incubate at 50°C for 1 hour to generate cDNA (SuperScript III, ThermoFisher Scientific). The HA1 sequence was divided into three fragments for sequencing. Seven Ns were added into the primers as the barcode for barcoded-subamplicon sequencing during the first round PCR (PrimeSTAR Max DNA Polymerase, Takara Bio, PCR condition was set according to the manufacturer). Equal amount of the products from the three fragments of each sample were purified (PureLink Quick Gel Extraction Kit, ThermoFisher Scientific) and mixed. 1.3×10^6^ copies of the PCR product from the mixture above were added into the second round PCR (KOD Hot Start DNA Polymerase, Novagen) to add the adapter for sequencing. The condition of the second round PCR was set according to the manufacture with an annealing temperature of 58°C for 25 cycles. The second round PCR products were then purified and submitted for sequencing on an SP line for 251 cycles from both ends of the fragments on a NovaSeq 6000 (250 nt, paired-end reads).

### Analysis of deep mutational scanning data

Sequencing data were processed as described before (Wang et al., 2021) to generate read counts for each amino acid substitution. The adaptors were trimmed, the FASTQ files were generated and demultiplexed with the bcl2fastq v2.20 Conversion Software (Illumina). Sequencing data was obtained in FASTQ format and analyzed using a custom Python snakemake pipeline (Mölder et al., 2021). First, UMIs were merged using python script with parameters: NNNNNNN 0.8 2, to obtain 5’-end paired UMIs with pattern ‘NNNNNNN’, at least two sequences with same UMIs, of which at least 80% are consensus sequence. Subsequently, primer sequences were trimmed using cutadapt (Martin, 2011), and then sequencing reads were renamed based on amplicon primer. Before variant calling and the fitness calculations, cleaned paired-end reads were merged by FLASH (Magoč and Salzberg, 2011) using parameters: -m 30 -M 70 -I. Finally, variants and normalized fitness values were calculated by python script as described previously (Wang et al., 2021). Briefly, merged paired-end sequences were firstly parsed by SeqIO module in BioPython (Cock et al., 2009) and then translated into protein sequences. Reads were filtered and removed if there is no amplicon tag or the sequence length was incorrect. Afterwards, variants of each residue were counted and normalized by amplicon to generate the frequency of each mutation. The enrichment ratio of each amino acid was calculated as follows:

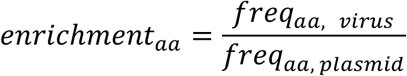

in which *freq*_*aa, virus*_ represents the frequency of a certain amino acid in the subamplicon in the output virus population after passage; *freq*_*aa, plasmid*_ represents the frequency of the amino acid in the subamplicon in the input plasmid.

The normalized fitness scores of individual amino acid substitutions were calculated based on the enrichment ratio of the amino acid normalized by the mean enrichment ratios of silent mutations and nonsense mutations (the average of enrichment ratio for all silent mutations and nonsense mutations calculated the same way as above):

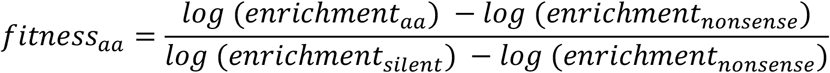

in which *enrichment*_*silent*_ and *enrichment*_*nonsense*_ are the mean of the enrichment ratio of the silent mutations and the nonsense mutations across HA1 for a given sample.

Custom python scripts for analyzing the deep mutational scanning data have been deposited to https://github.com/Wangyiquan95/HA1

### Monoclonal antibody selection

The saturated neutralization concentration was determined first for the selecting escape variant. Briefly, 10^7^ TCID50 of each virus were incubated with the monoclonal antibody with different concentrations in duplicate at 37°C for 30 min to equilibrate binding. Virus-antibody complexes were incubated with MDCK cells for 1 hour at 37°C. The cells were then washed by PBS and infection medium with the same concentration of antibody was added. The virus supernatants were then collected at 16 hpi and measured the titers by TCID50 assay. The saturated neutralization was decided by where the neutralization curve reached the plateau.

The virus stocks used for selection experiments were pooled from three independent rescues. The selection experiments were carried out as the same setting above with a saturate neutralization concentration escape variants were passaged if required and harvested at 16 hpi to obtain sufficient viral load (>10^4^ TCID50/mL) for high quality genome sequencing. No antibody treatment control groups were passaged in parallel. Viral RNA was extracted (QIAamp Viral RNA Kits, QIAGEN) from the supernatants and served as the template for RT-PCR (SuperScript III, ThermoFisher Scientific; Phusion High-Fidelity DNA Polymerase, ThermoFisher Scientific). For RT reaction, 10 μL of viral RNA and 1 μL MBTUni-12 primer (5′-ACG CGT GAT CAG CAA AAG CAG G-3′) was added into a 20 μL reaction. 10 μL cDNA template was amplified with MBTUni-12 and MBTUni-13 (5′-ACG CGT GAT CAG TAG AAA CAA GG-3′) primers and an annealing temperature at 57°C for 25 cycles. The input templates in PCR were normalized by the NP genome equivalents determined by qPCR. The PCR products were purified (PureLink Quick Gel Extraction Kit, ThermoFisher Scientific) and used to generate the shotgun library (KAPA HyperPrep Kits, Roche) and then sequenced one MiSeq flowcell for 251 cycles using a MiSeq 500-cycle sequencing kit version 2. The DeepSNV pipeline (Gerstung et al., 2012, 2014) was used to identify minor sequence variants. 2% minimum frequency threshold was set for variant calling.

### Biolayer interferometry (BLI)

The virus stocks used for BLI were purified by ultra-centrifugation as stated above. The protein concentration of the virus stock was determined by Bradford assay (Pierce(tm) Coomassie Plus (Bradford) Assay Kit, ThermoFisher Scientific). Streptavidin sensors (ForteBio) were coated with 500 nM 3’-SLN-PEG3-biotin (3’-Sialyllactosamine-PEG3-Biotin (Single Arm), Sussex Research). Equal concentrations of each virus were run on the BLI detection system (octet RED96e, ForteBio) for association for 300 seconds in the presence of 10 µM zanamivir to inhibit NA activity. Sensors were then incubated in PBS with zanamivir for 300 seconds for dissociation.

## Acknowledgements

This work has been generously supported by the National Institute of Allergy and Infectious Diseases of the National Institutes of Health under awards K22AI116588 and R01AI139246 to CBB, R00AI139445 and R01AI167910 to NCW, the Roy J. Carver Charitable Trust under award 17-4905 to CBB, and startup funds from the University of Illinois.

## Supplementary figures

**Figure S1:**
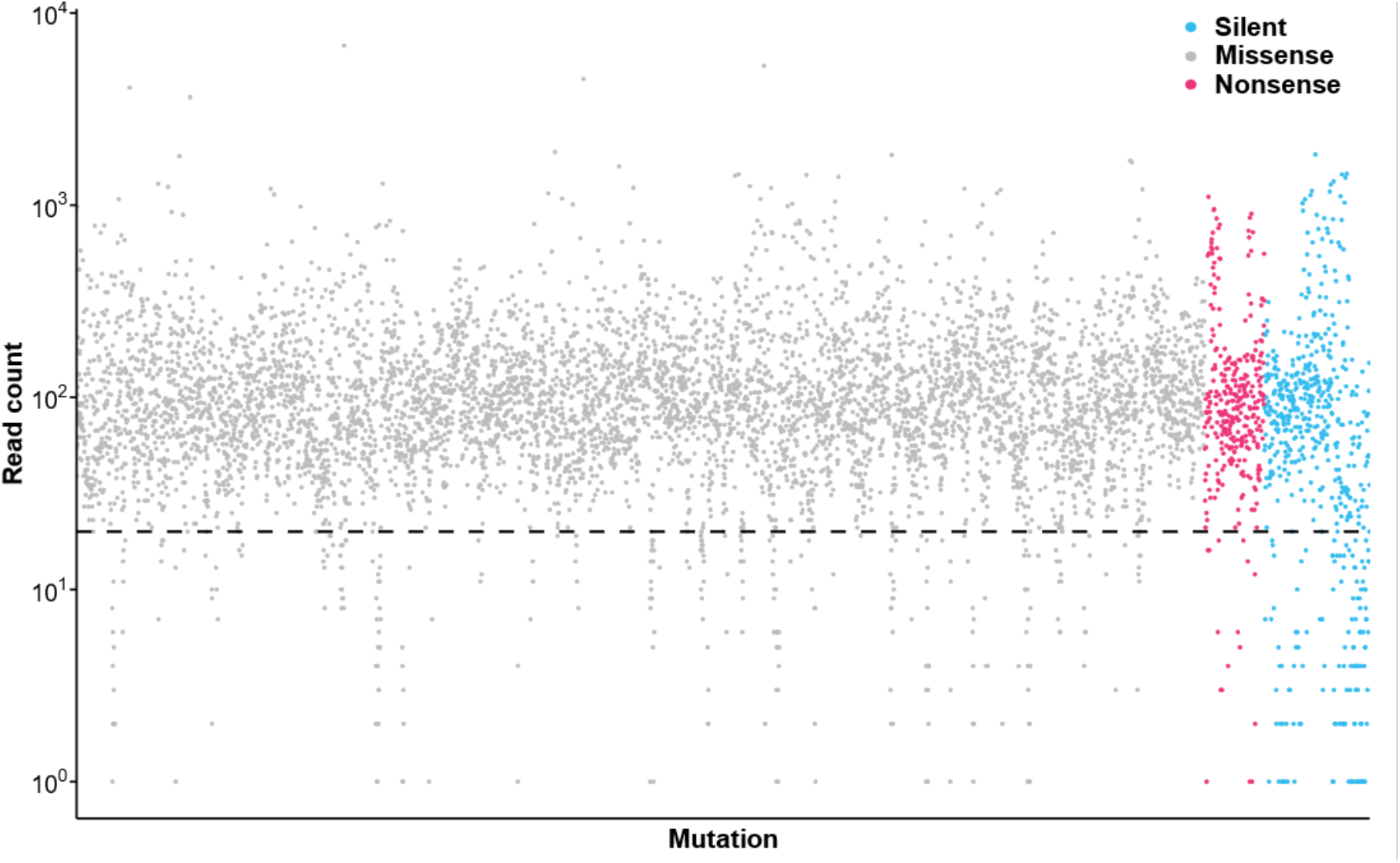
Mutation coverage in DMS plasmid library. Read count for every possible mutation (read count ≥ 1) in HA1. Dash line indicates the cutoff value (≥ 20) for downstream analysis (6303/6825). 168 mutations had read count = 0 in the plasmid library.

**Figure S2:**
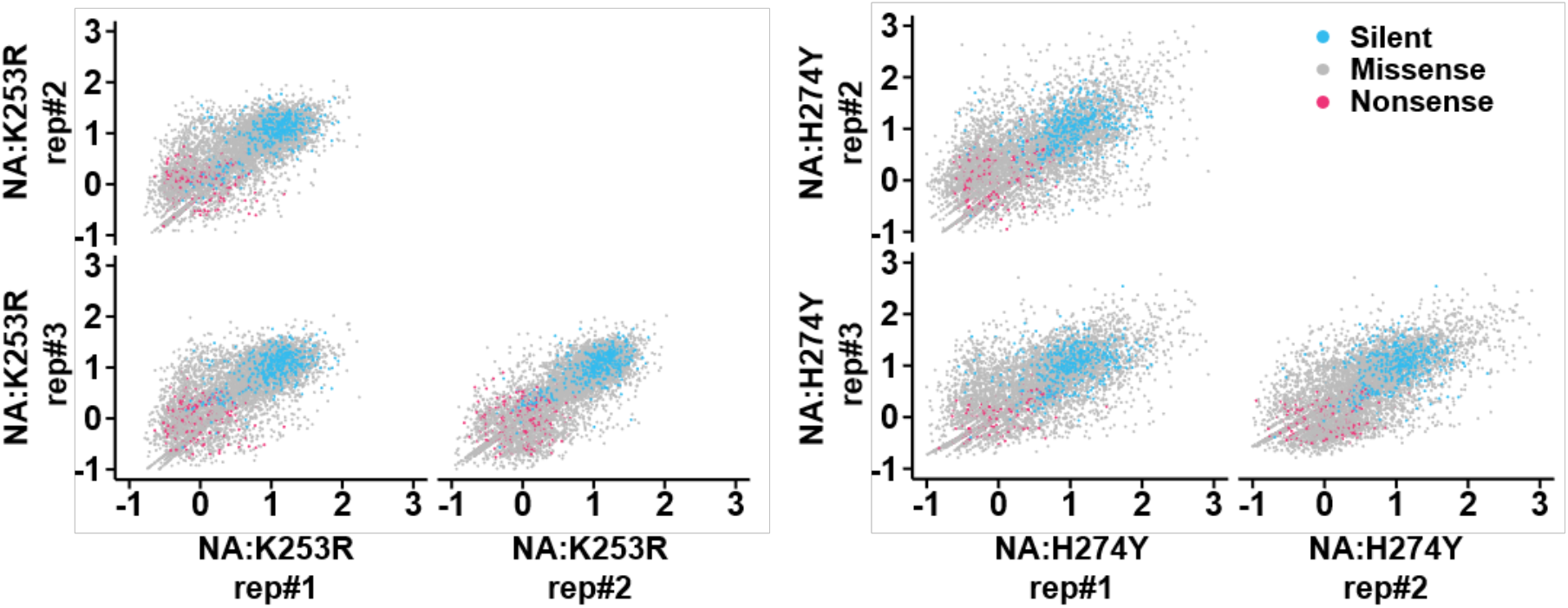
Correlation of normalized relative fitness score between replicates. Each dot represents the normalized relative fitness score of a specific substitution in the two indicated samples. Silent, missense, and nonsense substitutions are colored as indicated in the legend.

**Figure S3:**
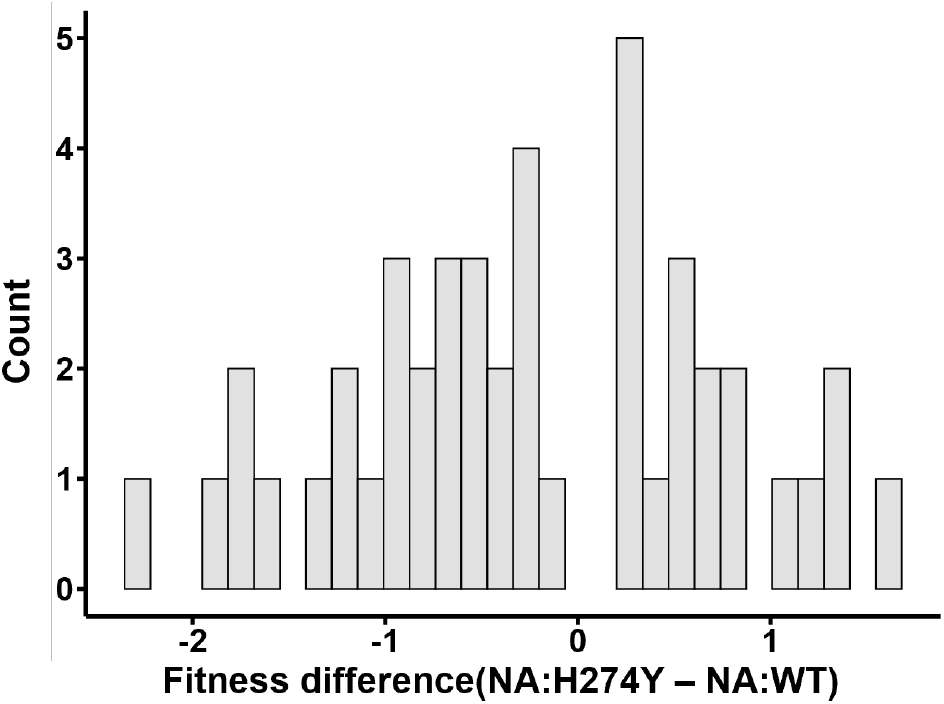
Fitness difference distribution between NA:H274Y and NA:WT. Data only show substitutions with differences with p < 0.01 based on t test.

**Figure S4:**
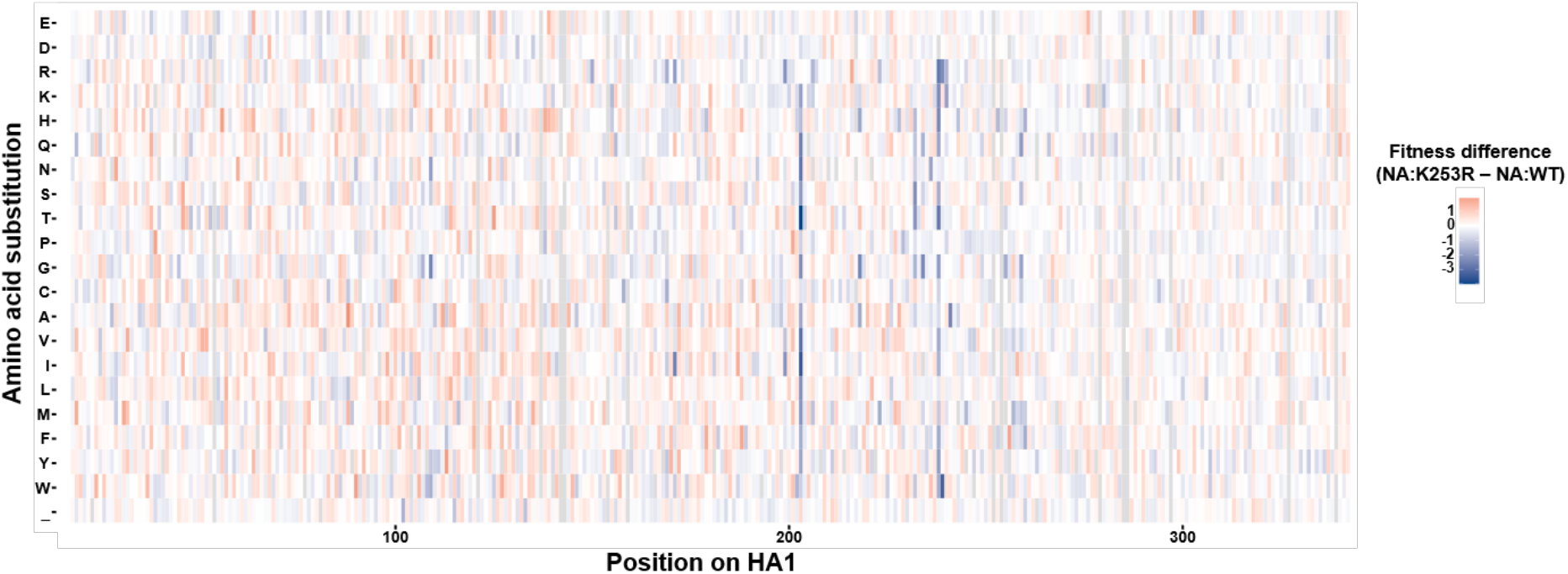
Fitness difference for each mutation between NA:H274Y and NA:WT. Normalized relative fitness score differences between NA:WT and NA:K253R for each substitution at each residue in HA1 as measured through DMS. Each value generated by subtracting the mean fitness score of 3 replicates for each genotype. Gray indicates that the mutation has the insufficient coverage in the plasmid library. The numbering was based on the H1 starting from the initiating methionine.

**Figure S5:**
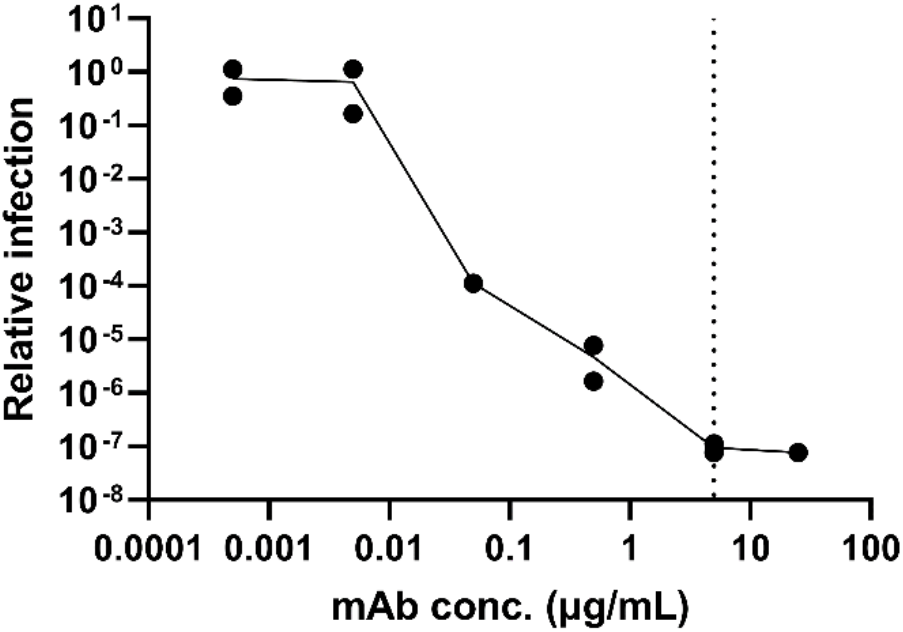
Saturated neutralization concentration of H36-26. 10^7^ TCID50 of NA:WT virus was neutralized by the given concentration of the antibody and infected a well of 6-well plate. The supernatant was collected 16 hours post infection. The output titer was measured by TCID50 assay and normalized by the titer of no antibody controls. The dash line indicates the concentration used in the selection experiment.

**Figure S6:**
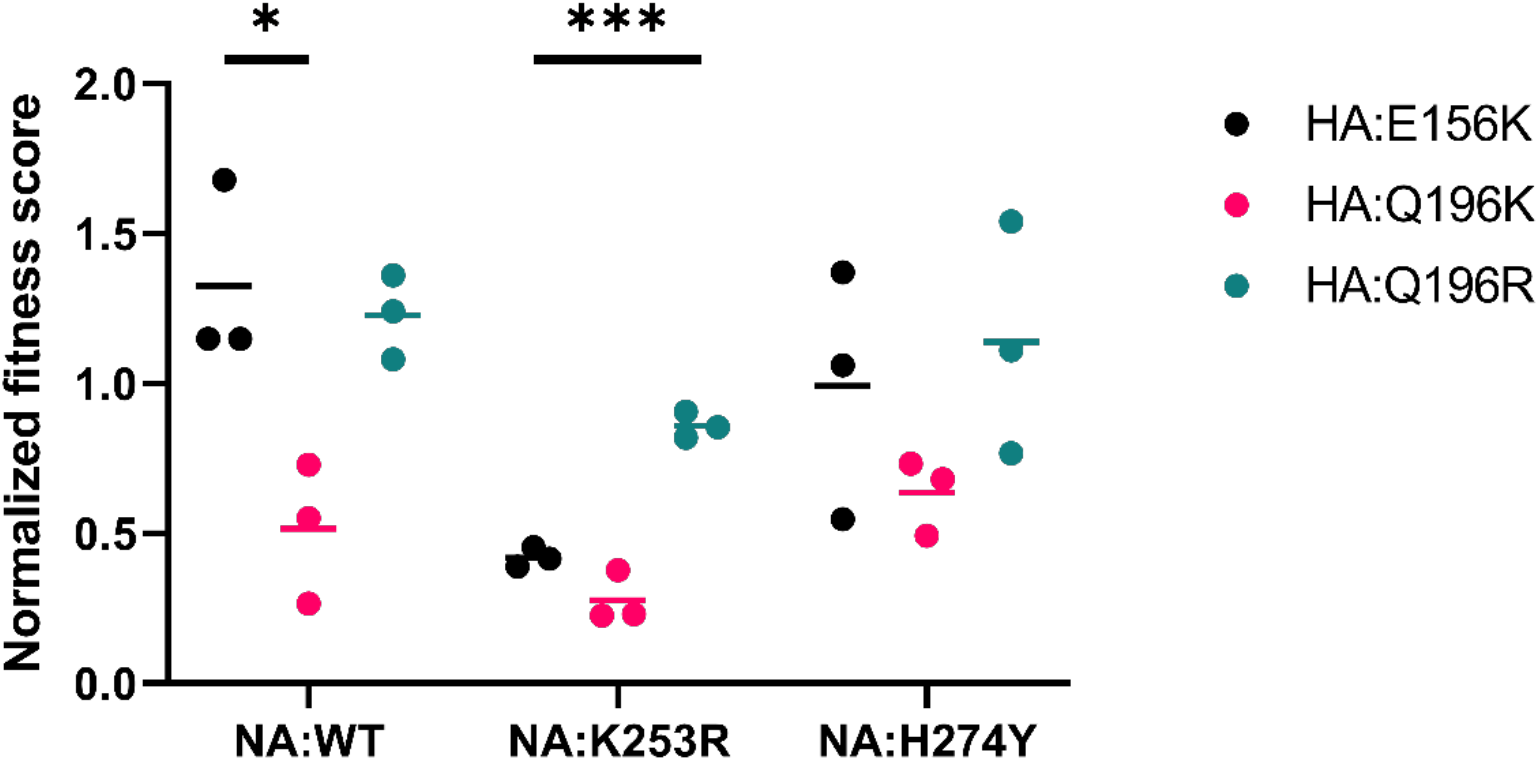
Fitness effect in DMS for H36-26 escape variants. The normalized relative fitness scores of the escape variants found in H36-26 antibody selection. *** indicates p < 0.01 and * indicates p < 0.05, p > 0.05 were not indicated in the graph, based on t tests.

**Figure S7:**
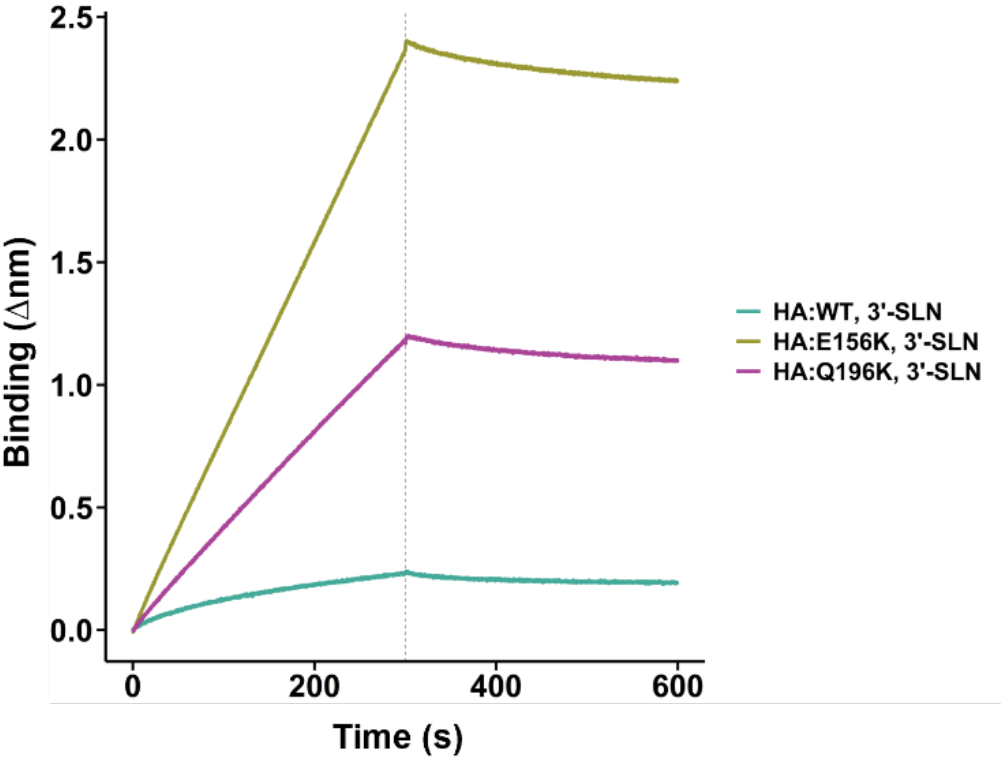
HA:E156K possesses higher receptor binding avidity. Binding kinetics to the receptor was measured by biolayer interferometry. The input was normalized by the protein concentration of the purified virion. Streptavidin sensors were coated with 3’-SLN-PEG3-biotin (3’-Sialyllactosamine-PEG3-Biotin (Single Arm). Separated by the dashed line, the first 300 seconds was the association period of the virion to the receptor, the next 300 seconds showed the dissociation period. 10 µM zanamivir was present during the assay to inhibit NA activity.

## Notes

### Competing Interest Statement

The authors have declared no competing interest.

